# Transfer Learning with False Negative Control Improves Polygenic Risk Prediction

**DOI:** 10.1101/2023.01.02.522532

**Authors:** X. Jessie Jeng, Yifei Hu, Jung-Ying Tzeng

## Abstract

Polygenic risk score (PRS) is a quantity that aggregates the effects of variants across the genome and estimates an individual’s genetic predisposition for a given trait. PRS analysis typically contains two input data sets: base data for effect size estimation and target data for individual-level prediction. Given the availability of large-scale base data, it becomes more common that the ancestral background of base and target data do not perfectly match. In this paper, we treat the GWAS summary information obtained in the base data as knowledge learned from a pre-trained model, and adopt a transfer learning framework to effectively leverage the knowledge learned from the base data that may or may not have similar ancestral background as the target samples to build prediction models for target individuals. Our proposed transfer learning framework consists of two main steps: (1) conducting false negative control (FNC) marginal screening to extract useful knowledge from the base data; and (2) performing joint model training to integrate the knowledge extracted from base data with the target training data for accurate trans-data prediction. This new approach can significantly enhance the computational and statistical efficiency of joint-model training, alleviate over-fitting, and facilitate more accurate trans-data prediction when heterogeneity level between target and base data sets is small or high.

**Author summary:** Polygenic risk score (PRS) can quantify the genetic predisposition for a trait. PRS construction typically contains two input datasets: base data for variant-effect estimation and target data for individual-level prediction. Given the availability of large-scale base data, it becomes common that the ancestral background of base and target data do not perfectly match. In this paper, we introduce a PRS method under a transfer learning framework to effectively leverage the knowledge learned from the base data that may or may not have similar background as the target samples to build prediction models for target individuals. Our method first utilizes a unique false-negative control strategy to extract useful information from base data while ensuring to retain a high proportion of true signals; it then applies the extracted information to re-train PRS models in a statistically and computationally efficient fashion. We use numerical studies based on simulated and real data to show that the proposed method can increase the accuracy and robustness of polygenic prediction across different ranges of heterogeneities between base and target data and sample sizes, reduce computational cost in model re-training, and result in more parsimonious models that can facilitate PRS interpretation and/or exploration of complex, non-additive PRS models.

## Introduction

Polygenic risk score (PRS), first proposed in [1], is a quantity that aggregates the effects of variants across the genome and estimates an individual’s genetic predisposition for a given trait. PRS can be used to explore the polygenetic architecture of a trait, to predict disease risk, to identify groups of individuals with substantially increased risks, and to study shared polygenic signals between different traits.

PRS is typically calculated as a weighted sum of trait-associated alleles, where the weights are the estimated effect sizes of the alleles on the trait. PRS analysis usually contains two input data sets: (i) base data, containing summary statistics of association tests from GWAS such as the estimated regression coefficients with the traits and their corresponding p-values, and (ii) target data, consisting of individual-level genome-wide genotypes data and traits used for prediction. Accumulating evidence suggests that for a variety of complex traits, variants selected based on a large ancestry-diversified base data are expected to harbor high fractions of causal variants carried by the target data, yet their effect sizes may not be the same across base and target data sets [2–4].

Because not all genetic variants influence the trait and because SNPs are correlated due to linkage disequilibrium (LD), two types of selection approaches have been adopted to construct PRS using GWAS summary statistics from the base data. The first type constructs PRS purely based on GWAS summary statistics from marginal model, i.e., removing correlated redundant SNPs via linkage disequilibrium (LD) pruning/clumping and computing the weighted allele sums only on those SNPs whose GWAS p-values are lower than a certain threshold (e.g., pruning/clumping + thresholding (C+T) [5], PRSice [6], and PRSice2 [7]). In these approaches, *p*-value threshold is usually set as a tuning parameter and optimized by data. The second type considers a joint additive model of all SNPs to estimate the effect sizes and to account for SNP LD, either by imposing lasso regularization (e.g., lassosum [8]) or prior distributions (e.g., LDpred [9], PRS-CS [10] and LDpred2 [11]). PRS based on joint modeling tend to have higher prediction accuracy [12] due to the increased accuracy in estimated SNP effects.

Given the utility of large-scale base data, it becomes more and more common to encounter the scenarios where the ancestral background of the base sample and target sample are not perfectly matched. For example, in the CoLaus/PsyCoLaus study [13, 14], the target sample is composed of Switzerland Caucasians while the base sample is composed of general European descents from US, central Europe, southern Europe and northern Europe. In multi-ethnic PRS prediction, the base sample tend to comprise European descents in large sample size as well as descents from other populations including the target population in moderate sample size. Consequently, although causal variants in the target data are expected to be included in a larger set of causal variants carried by the base data, the same variant may have different effect sizes in the base and target data sets.

To accommodate the potential ancestral heterogeneities between base sample and target sample, in this work, we treat the GWAS summary information obtained in the base data as knowledge learned from a pre-trained model, and adopt a transfer learning framework to leverage the knowledge learned from the relatively ancestry-diversified base data to build the PRS of target individuals. Our proposed trans-learning framework consists of two main steps: (1) conducting false-negative control (FNC) screening to extract useful knowledge from the base data; (2) conducting joint-model training to integrate the extracted knowledge with the target training data for accurate trans-data prediction. In (1), as SNP effects may differ across base and target datasets, marginal screening for promising SNPs should not just focus on strong base signals, but should have good capacity for weak base signals as well. Therefore, we use FNC to ensure the retention of a high proportion of strong and weak base signals and, at the same time, effectively exclude noise variants that are distinguishable from base signals. This is crucial for the second step of joint-model training, which would not be computationally efficient if not with a reduced set of “promising SNPs”, but would not be statistically powerful if not majority of causal SNPs are included in the reduced set. In (2), we use joint regression to train the target PRS model with individual-level target data on the reduced promising-SNP set, so as to ensure accurate SNP effect estimates that better reflect the underlying causal SNP architecture of the target sample that may be different from the base sample.

We use simulation to illustrate the performance of the proposed transfer learning procedure. In settings with various overlapping proportions of signals between base and target data sets, the new procedure leads to a better PRS model in terms of achieving a higher predictive *R*^2^ with a more parsimonious model. Application to the CoLaus/PsyCoLaus GWAS of lipid plasma indicates that besides being substantially faster than C+T, lassosum, and LDped methods, the new procedure can achieve the highest predictive *R*^2^ while retaining the least number of SNPs in the PRS model. A parsimonious PRS model that includes fewer SNPs whilst maintaining the same or higher predictive *R*^2^ can facilitate PRS interpretation and enable downstream analyses with more complex modeling techniques, such as incorporating SNP-SNP interactions in polygenic prediction models as discussed in [15, 16]. We also explore interactive models with the lipid prediction in the CoLaus/PsyCoLaus study.

## Materials and methods

### Formulation of trans-data signals

Let 𝒮 be the set of causal variants carried by the target data and 𝒮^+^ be the set of signal variants carried by the base data. The latter have non-null effects in the marginal models of GWAS. We consider heterogeneities between base data and target data from two sources: (a) the nested ethnicity relationship between base and target samples, and (b) the possible distortion of causal variants information from the pre-trained marginal models that have not accounted for LD. A general assumption on the trans-data information sharing can be

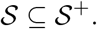

That is, the causal variants in the target data are included in the set of signal variants in the more ethnically diverse base data. 𝒮^+^ could also include “pseudo” signals, which have non-zero marginal effects because of their dependence with the causal variants in LD.

We do not impose the constraint that the effect size of a variant remains the same in different data sets. It is possible that the effect of an causal variant in 𝒮 gets diluted in the ethnically-mixed base data so that its effect is fairly weak in 𝒮^+^.

### Trans-data polygenic risk prediction

We propose a transfer learning polygenic prediction procedure which performs FNC marginal screening to effectively retain both strong and weak signals in 𝒮^+^ and applies the learned information from base data to identify causal variants 𝒮 in and improve PRS for target individuals. Different from the classical power analysis in hypothesis testing, FNC screening does not hinge on a pre-fixed control level of type I error (or some form of cumulative type I errors in multiple testing). Instead, FNC screening adapts to a user-specified level of false negative proportion (FNP) to facilitate more powerful discovery of weak signals while keeping out noise variants that are distinguishable from weak signals.

Figure 1 shows a flow chart of the transfer-learning procedure with FNC marginal screening and joint model training. In Step 1, we apply FNC marginal screening to the base data to extract useful knowledge as follows. First, we pre-fix a sequence of control levels, *ϵ*_1_, …, *ϵ*_*K*_, on FNP, which is the number of false negatives divided by the total number of signals | 𝒮^+^|. In this paper, we set *ϵ*_*k*_’s in grid values of {0.02, 0.04, …, 0.4}. Second, for a given *ϵ*_*k*_, we apply the FNC screening procedure of [17] on *p*-values of the base summary statistics and obtain a reduced SNP set 𝒟_*k*_, which is expected to contain a (1 *− ϵ*_*k*_) proportion of base signals in 𝒮^+^ as well as some noise SNPs. However, because FNC screening excludes noise SNPs that are distinguishable from the (1 *ϵ*_*k*_) proportion of signals in 𝒮^+^, each 𝒟_*k*_ can be much smaller than the full set of SNPs. More detailed descriptions of FNC screening is provided in the next subsection.

**Fig 1.**
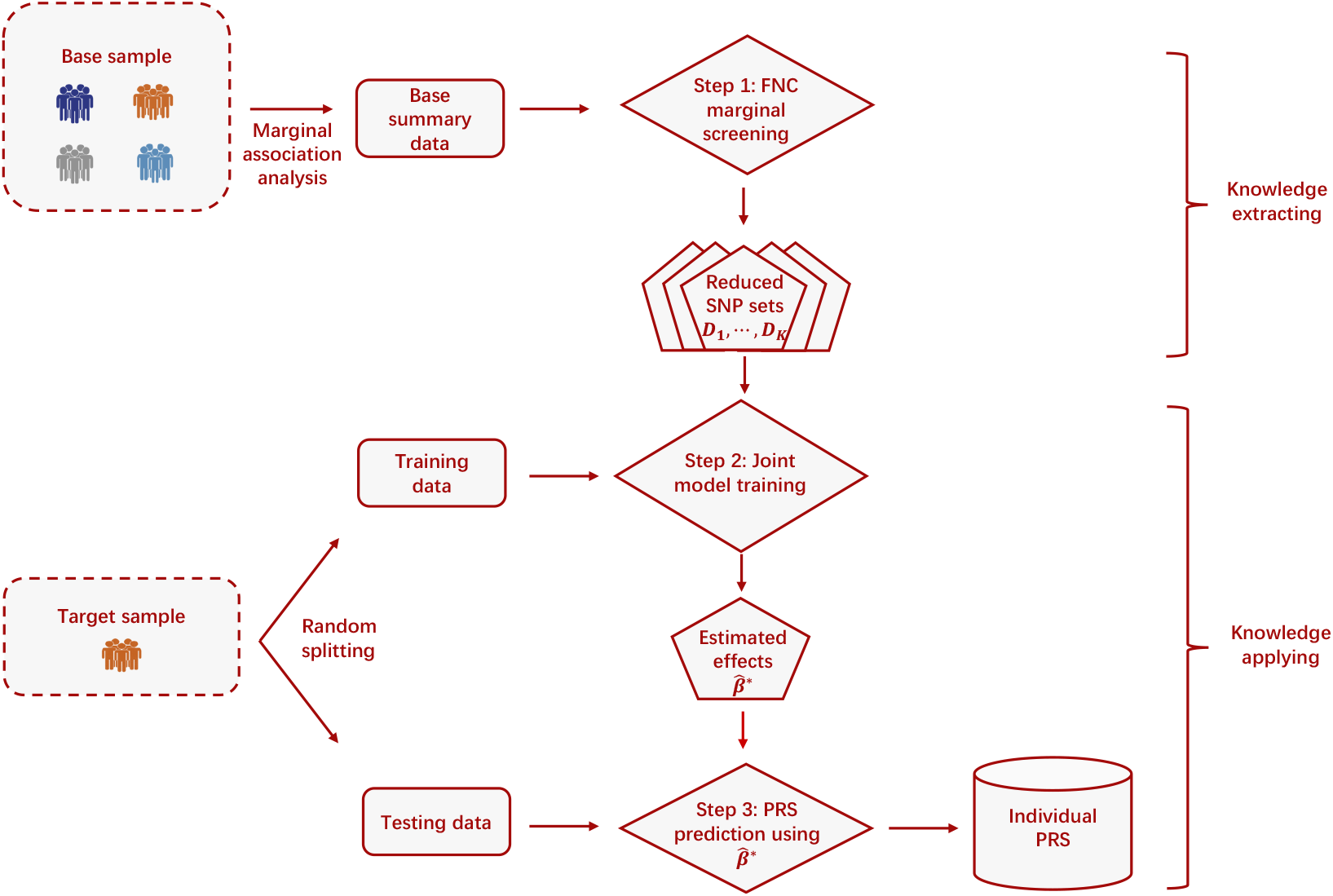
Overview of the PRS transfer learning procedure. Step 1: Given a pre-specified grid of control level for false negative proportion, {ϵ_1_, *…*, ϵ_*K*_} (e.g., {0.02, 0.04, *…*, 0.4}), apply false negative control (FNC) marginal screening to the base summary statistics and obtain *K* candidates of reduced SNP sets 𝒟_1_, *…*, 𝒟_*K*_, each of which retains a specific high proportion (i.e., 1 *− ϵ*_1_, *…*, 1 *− ϵ*_*K*_, respectively) of base signals. Step 2: For each reduced SNP set 𝒟_*k*_, train the joint prediction model using the target training sample. The joint model that yields the largest *R*^2^ is selected as the final model and the corresponding SNP effect estimates are denoted as 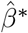. Step 3: Apply the estimated effects 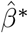 from the final model of Step 2 to the target testing sample to calculate individual PRS.

In Step 2, we equally and randomly divide the individual-level target sample into training and testing samples, and use the training sample and lasso regression to build the PRS model. Specifically, given a reduced SNP set 𝒟_*k*_, *k* = 1, …, *K*, we fit a joint prediction model using regularized regressions, select the regularization parameter using cross-validation or information criterion, and obtain the regularized estimates of SNP effects and compute the model *R*^2^ value. Among the *K* PRS models, the model yields the largest *R*^2^ value is selected as the final model, and the corresponding effect size estimates (denoted by 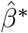) are used to construct PRS. In Step 2, working with the FNC reduced SNP set 𝒟_*k*_ instead of the entire SNP set enhances statistical power and computational efficiency of the joint-model training. In Step 3, we apply the estimated effects from the final model 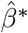 to the target testing sample to calculate individual PRS.

The proposed transfer learning procedure is general and can accommodate a spectrum of prediction models such as linear mixed effects models (LMM) [18] and penalized linear regression [19, 20] in Step 2. In this paper, we adopt the popular Lasso regression to obtain sparse effect estimates and utilize cross-validation to determine the hyper-parameters. We denote the whole procedure as FNC+Lasso and show the complete algorithm of FNC+Lasso in Algorithm 1 of S1 Appendix A. The computer code for FNC+Lasso, together with complete documentation is publicly available at Github: https://github.com/JessieJeng/FNC-Lasso.

### FNC marginal screening

The original FNC screening procedure was developed to retain a high proportion of signals via effective false negative control among arbitrarily dependent candidates [17]. FNC screening can be applied as follows. Given a sequence of *p*-values for all the SNPs in the base data: *p*_1_, …, *p*_*m*_, rank the SNPs by their *p*-values such at *p*_(1)_ *≤ p*_(2)_ *≤* … *≤ p*_(*m*)_. For each *j ∈ {*1, …, *m}*, calculate

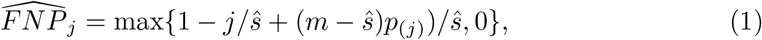

where ŝ is an estimate for | 𝒮^+^ |, the number signal variants in the base data. We implement the estimator introduced in [21] for dependent variants. Now, for a control level ϵ_*k*_, derive

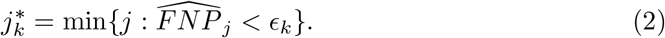

Then, construct the SNP set *𝒟*_*k*_ as the collection of SNPs whose *p*-values are less than or equal to 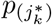. Repeat the above for each of ϵ_1_, …, ϵ_*K*_, and obtain the reduced SNP sets *D*_1_, …, *D*_*K*_.

The FNC screening procedure is based on direct estimation for the true FNP if only the top *j* ranked SNPs are selected. Rationale of the estimator in (1) is as follows. When the top *j* variants are selected, the total number of signal variants equals to the sum of true positives (*TP*_*j*_) and false negatives (*FN*_*j*_), i.e. |*𝒮*^+^| = *s* = *TP*_*j*_ + *FN*_*j*_. Then we have *FNP*_*j*_ = *FN*_*j*_*/s* = 1 − *TP*_*j*_*/s*, and we would need to estimate *s* and *TP*_*j*_ for each *j*.

FNC screening uses *ŝ* from [21] to estimate *s*, whose consistency has been proved under arbitrary covariance dependence. For *TP*_*j*_, note that selecting the top *j* variants incurs true positives and false positives with *j* = *TP*_*j*_ + *FP*_*j*_. Then *TP*_*j*_ = *j − FP*_*j*_. It is reasonable to use *E*(*FP*_*j*_) = (*m − ŝ*)*p*_(*j*)_ to approximate *FP*_*j*_ based on the null distribution of the p-values. Then *TP*_*j*_ can be approximated by *j −* (*m − ŝ*)*p*_(*j*)_ and, consequently, *FNP*_*j*_ can be approximated by 1 *− j/ŝ*+ (*m − ŝ*)*p*_(*j*)_*/ŝ* as in Equation (1).

Moreover, because the true FNP_*j*_ is non-increasing with respect to *j*, stopping at the first time 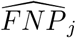 is less than ϵ_*k*_ can result in the smallest subset of variants with FNP controlled at ϵ_*k*_ with high probability. This step, as stated in Equation (2), can effectively exclude noise variants that are distinguishable from the (1 ϵ_*k*_) signals. More detailed justifications in theory and simulation of FNC screening can be found in [17].

## Results

### Simulation design

For the target data in simulation, we obtain the genotype data of Chromosome 21 from the CoLaus/PsyCoLaus study [13, 14]. After data pre-process described in S1B Text, the genotype data for Chromosome 21 has *m* = 5053 SNPs. We randomly select *n* individuals to form the target sample, resulting a genotype matrix *X* consisting of *m* columns and *n* rows. Let *β* be a *m*-dimensional vector of the allelic effects. We assume that *β* is a sparse vector with *β*_*j*_ ≠ 0 for *j ∈ 𝒮*, where 𝒮 is the set of causal variants for the target sample. The trait vector *Y* are simulated from *Y* = *Xβ* + *W*, where *W∼ N*_*n*_(0, *I*), *I* is an identity matrix. The simulated target sample is randomly split into training set and testing set with a 1 : 1 ratio. The training set is used to build PRS models using different methods. The testing set is used to evaluate the predictive performances of these methods.

The base summary data is generated by *Z ∼ N*_*m*_(*β*^+^, Σ*/n*_0_), where *n*_0_ represents the base sample size, Σ is the correlation matrix of all the *m* = 5053 SNPs, which is estimated from the CoLaus/PsyCoLaus sample. Such marginal effect model has been used in GWAS analysis in e.g. [22]. The mean vector *β*^+^ of effect sizes have 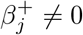 for *j ∈ 𝒮*^+^, and 0 otherwise, where *𝒮*^+^ is the set of base signals. In the simulation studies, we consider 3 sets of (*n*_0_, *n*), which are (4000, 2000), (4000, 1000) and (2000, 1000).

We control the overlap between base signals (^+^) and target signals () using a proportion parameter *δ* =|𝒮|*/* |𝒮^+^|, where represents the cardinality of a set. It is expected that the more diverse the base sample, the smaller the *δ*. In the simulation examples, we randomly selected 50 SNPs as causal variants in the target sample (i.e.,|*𝒮*| = 50) and set *δ* = 0.3, 0.5 or 0.7 representing low, median, and high overlaps between *𝒮* and *𝒮*^+^. The effect of each signal variant is generated independently from Uniform(0.05, 0.15) and separately for target and base data. The range of the uniform distribution is calibrated so that *R*^2^’s range from 5% to 30% for different methods across different scenarios.

We simulate 100 replicates under each simulation scenario, and evaluate the predictive performance, model fitting and model parsimony for the final PRS model. Specifically, we examine two metrics: (1) *R*^2^, which quantifies the outcome variation explained by the PRS on the target testing set; and (2) Akaike Information Criterion (AIC), which evaluates how well the PRS model fits the outcome data and has been used to evaluate polygenic prediction in [23] and [24]. AIC is calculated by *AIC* = 2*q* + *n*_*test*_ *·* log(*RSS/n*_*test*_)), where *q* is the number of SNPs retained in the PRS model, *n*_*test*_ is the sample size of the target testing set, and *RSS* is the residual sum of square (RSS) calculated on the target testing set. As an ancillary metric, we also report the number of SNPs included in the final PRS model.

### Baseline Methods

We benchmark the proposed FNC+Lasso transfer learning method against five representative methods for PRS analysis, which are Clumping+Thresholding [CT; [5]], lassosum [8], LDpred [9, 11], Lasso [25], and SIS+Lasso. The first three methods utilize base summary data to estimate the effect sizes, and were implemented using R package `bingsnpr` [26] in our numerical studies. In contrast, Lasso only uses the individual-level target data, see for example [20]. The last method, SIS+Lasso, can be viewed as a naive transfer learning procedure that also includes dimension reduction and joint-model training. Instead of using FNC screening and searching adaptively for the optimal reduced SNP set, SIS+Lasso applies sure independence screening (SIS) [27] to the base summary statistics to select the top SNPs based on the training sample size, and then performs Lasso regression on the selected SNPs with the target training data. It is well-known that SIS can be applied to marginal summary statistics to quickly reduce data dimension. We include SIS+Lasso in the baseline methods to evaluate potential advantages of using FNC screening in the transfer learning framework.

### Simulation Results

Figure 2 shows the boxplots of *R*^2^ from 100 replications, for (*n*_0_, *n*) = (4000, 2000) (top row), (4000, 1000) (middle row) and (2000, 1000) (bottom row). In each replication, the target sample is randomly and equally divided into training and testing sets. In all scenarios, we observed that FNC+Lasso tend to give the highest *R*^2^ among all methods. On one hand, methods using a trans-learning framework (i.e., FNC+Lasso and SIS+Lasso) generally outperform other baseline methods. The only exception is when the training sample size is relatively small (e.g., *n* = 1000) and the overlap between base and target signals is high (e.g., *δ* = 0.7), where CT can perform comparable to or better than SIS+Lasso. On the other hand, FNC+Lasso, which aims to reserve an optimal high proportion of true signals during screening, consistently yield better *R*^2^ than SIS+Lasso and other baseline methods.

**Fig 2.**
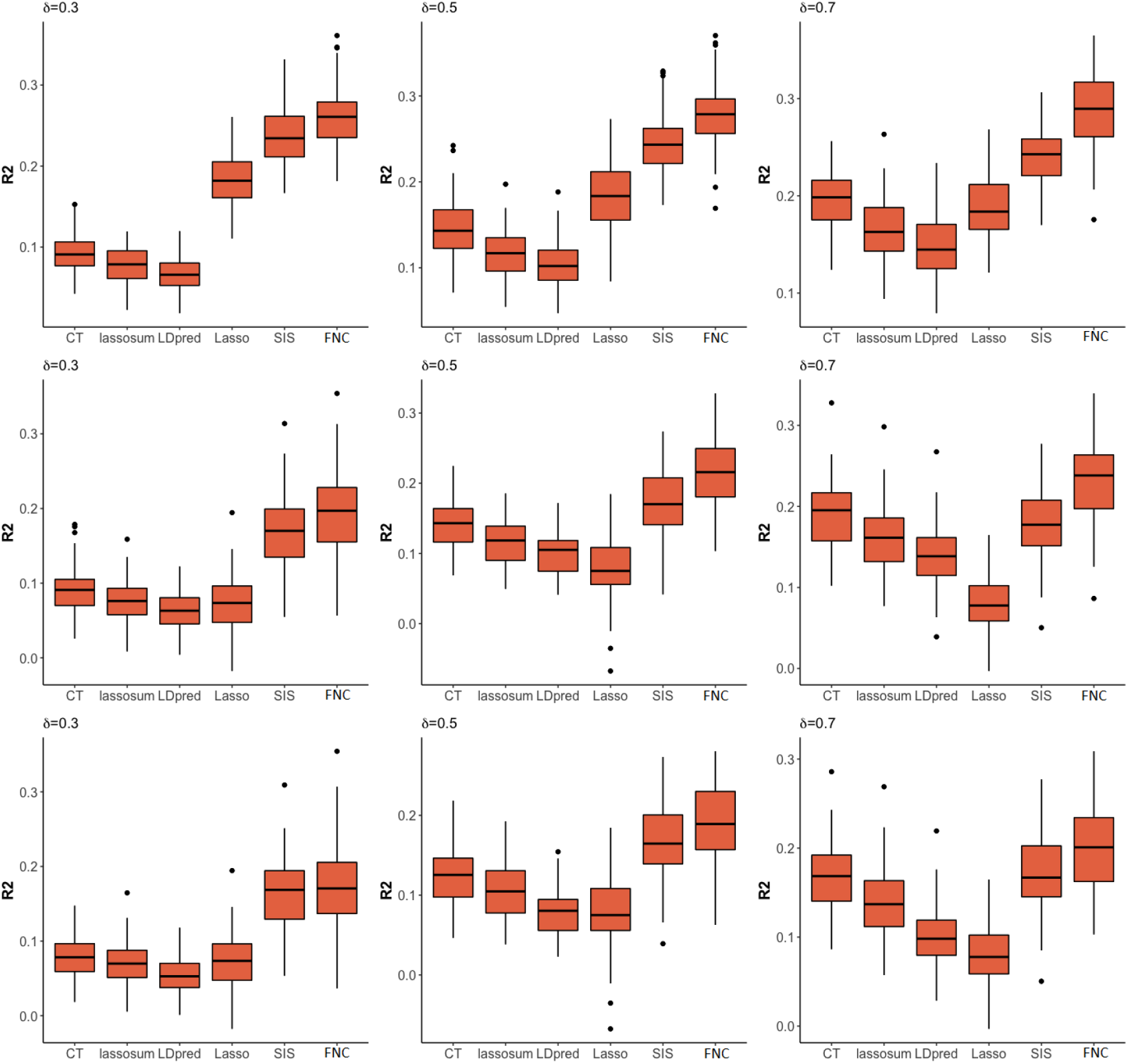
Results of *R*^2^ for prediction accuracy of different PRS methods. The results are based on 100 simulation replicates under different sample-size combinations of base data (*n*_0_) and target data (*n*) and different overlap proportion between base signals and target signals (*δ*). From the top to bottom rows are for (*n*_0_, *n*) = (4000, 2000), (4000, 1000) and (2000, 1000), respectively. In each replication, the target sample is randomly and equally divided into training and testing sets. From the left to right columns are for *δ* = 0.3, 0.5, and 0.7, respectively. Methods considered include Clumping+Thresholding (CT), lassosum, LDpred, Lasso, SIS+Lasso (SIS), and FNC+Lasso (FNC).

The relative performances of the non-trans-learning baseline methods are sensitive to the target sample size *n* and the overlap proportion *δ*. Specifically, when target sample size is relatively large (e.g., *n* = 2000; top row of Figure 2), Lasso tends to have higher *R*^2^ than those methods relying on the effect sizes estimated from base data, i.e., CT, lassosum, and LDpred. The gap is more obvious when the overlap proportion *δ* is small. The results suggest that re-estimating SNP effects using target training data (i.e., Lasso) can improve predictive performance when the target sample size is sufficiently large, and that the gain is more substantial when the base signals and target signals are less homogeneous (i.e., smaller *δ*).

In contrast, when target sample size is relatively small (e.g., *n* = 1000; middle and bottom rows of Figure 2), CT, lassosum, and LDpred generally outperform Lasso. The gap is more obvious when the overlap proportion *δ* becomes larger. The results suggest that when target sample size *n* (and hence the target training sample size *n*_*train*_) is not sufficiently large, it becomes inefficient to re-estimate SNP effects using target training data, even when the base signals and target signals become more heterogeneous (e.g., overlap proportion *δ* = 0.3 or 0.5). When the overlap between base and target signal sets increases, relying on the base effect estimates, which are obtained on a larger sample size *n*_0_, can enhance predictive *R*^2^ (e.g., CT has much higher *R*^2^ than Lasso and even similar *R*^2^ as SIS+Lasso when *δ* = 0.7 in the middle and bottom rows). We also observe that the results of (*n*_0_, *n*) = (4000, 1000) and (2000, 1000) are very similar, implying that the base sample size *n*_0_ has less impact on relative performance of the baseline methods.

Table 1 shows AIC and the number of SNPs included in the final PRS model. A smaller AIC value indicates a better model fitting in simplicity and accuracy. In all the scenarios, the proposed FNC+Lasso achieves the lowest AIC value and selects the least amount of SNPs among all PRS methods. Across various base and target sample combinations, (*n*_0_, *n*), SIS+Lasso tends to have the second smallest AIC when the overlap proportion is not low (e.g., *δ* = 0.3 and 0.5); when *δ* is high (e.g., 0.7), CT yield PRS models with smaller AIC and less SNPs than SIS+Lasso.

**Table 1.**
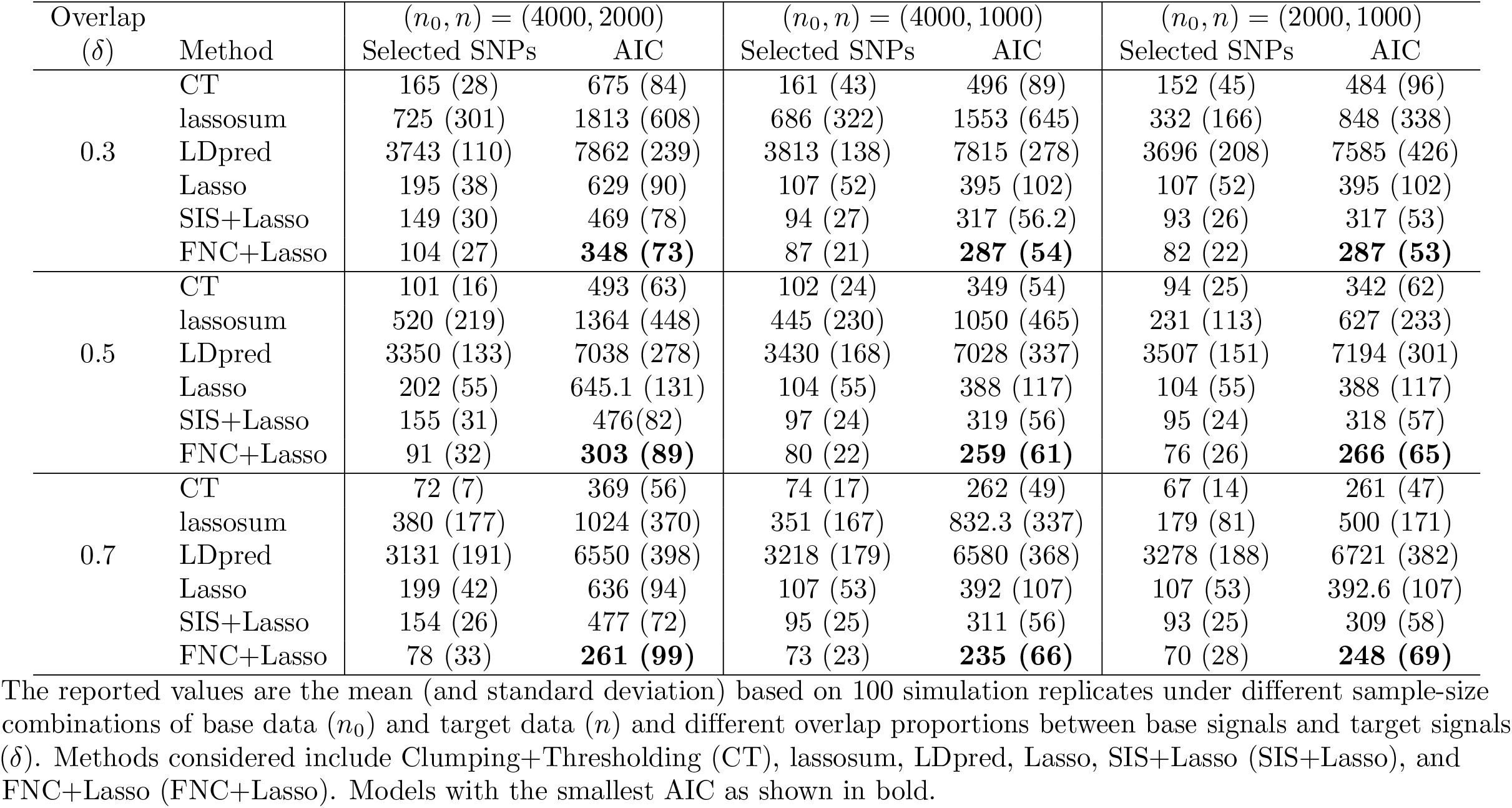
Results of Akaike Information Criterion (AIC) and number of SNPs in the final PRS model of different methods for assessing model fit and parsimony.

### Real data applications

We applied the proposed FNC+Lasso method and the baseline methods to construct the PRSs for plasma lipids, including total cholesterol (CHOL), triglycerides (TRIG), high-density lipoprotein (HDL) and low-density lipoprotein (LDL). Plasma lipids are risk factors for cardiovascular diseases, a leading cause of death worldwide [28], and CHOL, TRIG, HDL and LDL are effective biomarkers and risk factors for heart attack and heart diseases. At present, over 1,900 common SNPs have been shown to be associated with CHOL with different significant levels and the corresponding number for TRIG is 4,002 [29]. However, majority of these variants confer small risk individually and have limited predictive power for lipid traits [30], although the variance proportions explained by PRS for Whites and Hispanics tend to be higher than other populations [31].

Our target data is obtained from the CoLaus/PsyCoLaus GWAS study of the Cohort Lausanne, Switzerland [13, 14], consisting of 5,247 Switzerland Caucasians individuals. We acquired the base data from [32] and [33], which includes samples of general European descents from US, central Europe (i.e., Australia, Denmark, France Germany, Netherlands, Switzerland, UK), southern Europe (Italy) and northern Europe (i.e., Iceland, Finland, and Sweden). The base sample has broader ancestral background and larger sample sizes (i.e., 94,595, 100,184, 99,900 and 95,454 for CHOL, TRIG, LDL and HDL, respectively). We pre-process the genotype data as detailed in S1 Appendix B and matched SNPs between the base and target data. The number of matched SNPs for CHOL, TRIG, LDL and HDL are 268,961, 266,952, 268,939 and 268,939, respectively.

To construct PRSs, we randomly split the target sample into training and testing sets, each containing half of the target sample. We apply the six methods, i.e., the 5 baseline methods (CT, lassosum, LDpred, Lasso and SIS+Lasso) and the proposed FNC+Lasso. In all real data analyses, we include the top 10 principal components for population substructure, sex and age as covariates. We optimize the tuning parameters and obtain the effect size estimates for the final PRS model in training set, and calculate the prediction *R*^2^ of the final PRS model in testing set. To account for the variations involved in the random data splits, we repeat the process using 50 different splitting of training and testing sets. In this part, we use the R package *bigstatr* for Lasso to ensure computational efficiency with whole genome SNPs.

Table 2 shows the *R*^2^, AIC and number of selected SNPs of different methods. Across all four traits, the proposed FNC+Lasso yields more robust performance in *R*^2^ and AIC by yielding the best or comparable values to the best-performing methods, while the best-performing methods can vary depending on traits, *R*^2^ and AIC. For example, for CHOL, lassosum has the smallest AIC but lower *R*^2^ (4.8%) than SIS+Lasso (5.7%) and FNC+lasso (6.3%). For TRIG, FNC+Lasso has the higest *R*^2^ and lowest AIC. For LDL, LDpred has the highest *R*^2^ but a much larger AIC than Lasso, SIS+Lasso and FNC+Lasso. For HDL, CT has the highest *R*^2^ and lowest AIC. We also observe that although giving reasonable AICs, the classical Lasso has the lowest *R*^2^ for all four traits, suggesting that re-training a PRS model using Lasso on all SNPs with target sample does not yield ideal predictive power. Incorporating a screening procedure using base data (i.e., SIS+Lasso and FNC+Lasso) can substantially enhance the predictive *R*^2^ of Lasso. However, only FNC+Lasso can consistently yield a comparable or higher *R*^2^ than the best-performing methods, which is FNC+Lasso (for CHOL and TRIG), LDpred (for LDL), and CT (for HDL).

**Table 2.** Prediction *R*^2^, number of selected SNPs, AIC, and computing time of different PRS methods for the CoLaus/PsyCoLaus analysis of four lipid traits.

| Trait | Method | Predictive $R^2$ | Selected SNPs | AIC | Computing time (sec) |
| --- | --- | --- | --- | --- | --- |
| CHOL | CT | 4.5% (0.4%) | 506 (618) | 1134 (1232) | 753 (13) |
|  | lassosum | 4.8% (0.6%) | 5 (1) | **124 (56)** | 495 (4) |
|  | LDpred | 4.7% (0.5%) | 219930 (58282) | 205687 (11652) | 1860 (7) |
|  | Lasso | 3.2% (0.1%) | 208 (169) | 572 (338) | 1232 (1554) |
|  | SIS+Lasso | 5.7% (0.4%) | 406 (96) | 901 (185) | **5 (1)** |
|  | FNC+Lasso | **6.3% (0.7%)** | 127 (38) | 324 (88) | 129 (11) |
| TRIG | CT | 5.6% (0.4%) | 52403 (47562) | 105693 (95099) | 739 (55) |
|  | lassosum | 5.2% (0.5%) | 21921 (61078) | 44774 (122156) | 589 (23) |
|  | LDpred | 5.7% (0.5%) | 113050 (6254) | 226987 (12467) | 1879 (5) |
|  | Lasso | 3.6% (0.3%) | 235 (180) | 1411 (582) | 126 (21) |
|  | SIS+Lasso | 5.7% (0.6%) | 315 (146) | 1515 (529) | **10 (4)** |
|  | FNC+Lasso | **7.1% (0.7%)** | 93 (15) | **1033 (359)** | 186 (22) |
| LDL | CT | 4.1% (0.8%) | 30154 (40143) | 59815 (80277) | 647 (44) |
|  | lassosum | 4.0% (0.6%) | 16189 (17905) | 31875 (35802) | 655 (13) |
|  | LDpred | **4.4% (0.6%)** | 7912 (1221) | 15323 (2448) | 1828 (7) |
|  | Lasso | 1.9% (0.2%) | 219 (128) | 5 (274) | 182 (64) |
|  | SIS+Lasso | 2.6% (0.2%) | 135 (68) | -184 (169) | **8 (4)** |
|  | FNC+Lasso | 3.5% (0.4%) | 66 (15) | **-346 (74)** | 138 (22) |
| HDL | CT | **23.2% (2.1%)** | 79 (54) | **-4864 (132)** | 589 (185) |
|  | lassosum | 20% (0.1%) | 9041 (13941) | 13141 (27872) | 592 (8) |
|  | LDpred | 18.9% (1.0%) | 8367 (101) | 11836 (58) | 1842 (22) |
|  | Lasso | 17.8% (0.8%) | 362 (247) | -4121 (469) | 1947 (895) |
|  | SIS+Lasso | 21.5% (1.3%) | 445 (93) | -4077 (158) | **4 (1)** |
|  | FNC+Lasso | 22.7% (1.5%) | 207 (74) | -4590 (161) | 103 (15) |

The reported values are the mean (and standard deviation) from 50 random splits for each of the four lipid traits, including total cholesterol (CHOL), triglycerides (TRIG), high-density lipoprotein (HDL) and low-density lipoprotein (LDL). Methods considered include Clumping+Thresholding (CT), lassoSum, LDpred, Lasso, SIS+Lasso (SIS+Lasso), and FNC+Lasso (FNC+Lasso). Best performing results across different methods are shown in bold.

From the number of selected SNPs in each PRS model, we find CT, lassosum and LDpred tend to select tremendous number of SNPs for prediction. Such results are not unexpected for methods utilizing effect sizes estimated from the base data that are sampled from a broader population. The methods relying on Lasso (i.e., Lasso, SIS+Lasso and FNC+Lasso) to estimate effect sizes from target data contain much smaller set of selected SNPs. We note that the proposed FNC+Lasso tends to use the less number of SNPs while achieving a higher predictive *R*^2^, gives the smallest or the second best AICs, and has a more robust performance across different traits.

The parsimonious modeling of FNC+Lasso could foster interpretability and facilitate more complex modeling in downstream analyses, such as incorporating SNP-SNP interactions in polygenic prediction models. Here we explore pairwise interactions of the SNPs selected in FNC+Lasso on the lipid traits. To avoid potential overestimation of the interaction effects, we fit an linear regression model that satisfies strong hierarchy (i.e., whenever a SNP-SNP interaction is estimated to be non-zero by Lasso, the main effects of both SNPs are also included in the model) so that the interaction effects are not identified due to their collinearity with the omitted main effects. This procedure utilizes hierarchical group-lasso regulation introduced in [34] and can be applied using the R package *glinternet*. We investigate the predictive *R*^2^ performance of the interaction PRS model based on FNC-Lasso selected SNPs (denoted as FNC-INT) for the four traits. The results in Figure 3 show that incorporating SNP-SNP interaction terms can further increase the predictive performance for CHOL and LDL, although interactions do not help in TRIG and HDL. Recent studies have gain valuable insights into the functional consequences of some gene-gene interactions on LDL through systematic experiments that simultaneously knockdown candidate gene pairs in cultured HeLa cells [35]. Further follow-up studies could relate our identified SNP-SNP pairs to candidate genes and gene pairs through eQTL analysis and evaluate their roles in biological pathways of lipid traits.

**Fig 3.**
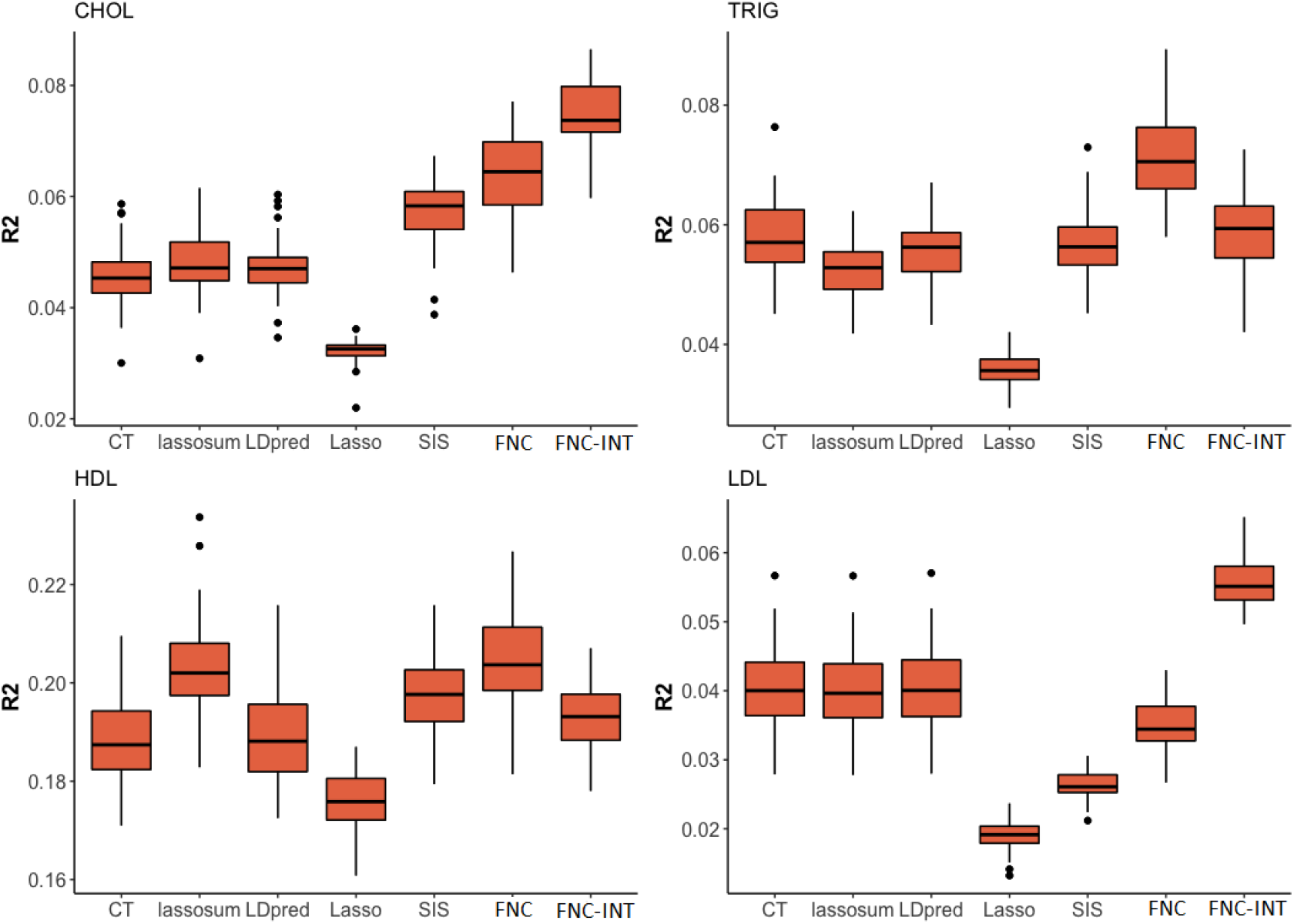
*R*^2^ values for prediction accuracy of different PRS methods. Methods considered include Clumping+Thresholding (CT), lassosum, LDpred, Lasso, SIS+Lasso (SIS), FNC+Lasso (FNC), and interaction model based on FNC+Lasso identified SNPs (FNC-INT) for four lipid traits.

Table 2 also shows that the proposed FNC+Lasso is computational efficient. Although FNC+Lasso involves a grid search to determine two hyper-parameters (FN proportion ϵ_*k*_ and the shrinkage parameter of Lasso), the required computing time is still substantially less than CT, lassosum, LDpred, and Lasso, e.g., FNC+Lasso can be 4-14 times faster than these baseline methods except for SIS+Lasso.

### Conclusion and discussion

Inspired by the idea of transfer learning, the proposed FNC+Lasso procedure aims to leverage the base data information for target data prediction modeling, where the target data may or may not have similar ancestral background as the base data. FNC+Lasso utilizes a unique false negative control strategy to extract useful information from the base data, and applies the extracted information to re-train the PRS models using target data in a statistically and computationally efficient fashion. The benefits are multi-fold: we can obtain more accurate polygenic prediction, reduce cost in model training, alleviate over-fitting, and facilitate model interpretation and complex modeling in follow-up analyses. The proposed PRS transfer learning framework can accommodate continuous and categorical traits by applying linear or logistic models in the joint modeling step. It is also applicable to the target data with only summary statistics available by implementing, e.g., lassosum in the joint modeling step.

We measure the overlap of signals between target and base data by a proportion parameter *δ*, which is unknown in real applications and can vary from trait to trait. It is expected that when signal overlap is low (i.e., heterogeneous signal patterns between base and target data), re-training PRS models with sufficient target samples can achieve better predictive performance. When signal overlap is high (i.e., similar signal patterns between base and target data), utilizing effect estimates from base data can ensure higher predictive accuracy, especially when target samples are of limited size. We observe such trends in our numerical studies when examining the relative performance of baseline methods. Our numerical studies also show that the proposed FNC+Lasso adapts to signal overlap level — FNC+Lasso ensures effective PRS joint-model training even with limited target samples, and results in a more robust and better predictive performance than the baseline methods across a range of *δ* values and different base/target sample sizes. As more samples are available for target and base data sets, the transfer learning method would benefit from integrating knowledge from both data sources with arbitrary signal overlaps and provide more accurate polygenic prediction.

The PRS models of FNC+Lasso tend to be more parsimonious compared to baseline models while maintaining a comparable or higher *R*^2^. A parsimonious PRS model that consists of key SNPs and excludes noise SNPs could aid the interpretablity of PRS and help to inform underlying molecular and functional basis, such as by examining the SNPs involved in the PRS model as well as those SNPs in LD with them to identify biologically important variants or to identify hub genes that capture larger polygenicity [36]. By focusing on the SNPs in a parsimonious PRS model, it also allows further exploration of non-additive PRS models, such as the SNP-SNP interaction PRS model considered in the CoLaus/PsyCoLaus data analysis. PRS models incorporating epistasis have been challenging because existing PRS models tend to be comprised of a large number of SNPs, and the corresponding interactions quickly leads to a prohibitive number of predictors in the model. While machine-learning PRS methods such as random forest have been proposed to account for unknown epistasis [37], it remains an open question how the base data information can be incorporated into these learning approaches. Complex models based on FNC+Lasso SNPs may offer an alternative for constructing non-additive PRS.

## Supporting information

**S1 Appendix. FNC+Lasso algorithm and preprocess of the CoLaus/PsyCoLaus GWAS data**. Section A shows the FNC+Lasso algorithm. Section B describes the preprocess of the CoLaus/PsyCoLaus GWAS data.

## Acknowledgments

The authors thank Dr. Peter Vollenweider, Dr. Gerard Waeber, Dr. Julien Vaucher and Dr. Martin Preisig, PIs of the CoLaus/PsyCoLaus study, and collaborators at GSK for providing the CoLaus/PsyCoLaus phenotype and genotype data.

